# Island Biogeography Theory of coevolution in pollination networks

**DOI:** 10.1101/2025.10.22.684063

**Authors:** Irina Birskis-Barros, Justin D. Yeakel

## Abstract

Island ecosystems have been pivotal in understanding community assembly and biodiversity, from the competing roles of colonization and extinction, to the influence of spatial structure, species interactions, and evolutionary processes. Network theory has helped our understanding of how ecological interactions shape island biogeography dynamics, using empirical and theoretical studies to explore species-rich island community structures. Mutualistic networks on islands differ from those on the mainland by supporting fewer species and more super-generalists, resulting in a more nested community structure. Coevolution, the reciprocal adaptation between interacting species, can shape traits and interactions within these networks, influencing their assembly. Here we explore how colonization, extinction, and coevolution can intersect to shape species traits and the structure of mutualistic networks. Using a stochastic dynamic model, we integrate Island Biogeography Theory with coevolutionary dynamics, examining pollination networks to understand trait-matching in evolving communities. Our results show that highly nested and connected mainland communities contribute to greater instability on islands, particularly under intermediate extinction rates. While super-generalist species persist longer, their persistence does not greatly exceed those species with fewer interactions beyond a threshold, which itself varies with the extinction rate. Additionally, coevolution leads to greater trait-similarity within communities on islands. Our findings highlight the critical role of coevolution in shaping island communities, especially on small islands where the extinction rate is expected to be elevated.

## 1 Introduction

Island ecosystems have long played a critical role in our understanding of community assembly, offering insights into the dynamic processes shaping biodiversity. MacArthur and Wilson [1] proposed the influential Theory of Island Biogeography, describing how colonization and extinction processes can influence island biodiversity over time, leading to species turnover and the eventual attainment of equilibrial community richness. Many empirical studies have demonstrated where these theoretical insights align with observations of island community assembly, as well as inherent limitations. For example, factors such as archipelago structure and history [2, 3], speciation [4], and the structure and dynamics of species interactions [5] are critical components that also shape island communities, expanding beyond the original scope of the theory.

Species interactions not only constrain the distribution of species, but can also facilitate species coexistence [6, 7]. The effects of multiple species interactions and the complex structure of ecological systems can be investigated using tools from network theory [*e*.*g*. 8–10]. In this sense, ecological communities can be represented as networks in which species represent nodes of the network and if they interact, are connected by edges, or links. An edge connecting two species can represent many different types of relationships, but is often used to denote energetic or biomass flow transferred between a consumer and resource species engaged in a trophic interaction [11, 12]. Advances in network theory have deepened our understanding of how ecological interactions shape island biogeography dynamics, with both empirical and theoretical studies applying these concepts to explore the structure of species-rich island communities [13–16]. For example, Gravel et al. [5] demonstrated the importance of integrating food web structure into the classical Island Biogeography Theory model to improve predictions of real community composition on islands. By incorporating the assumption that a consumer species must have at least one prey species present to colonize the island and that losing all prey results in extinction, they showed that the structure of the food web can play a crucial role in determining the species richness of island communities [5].

While trophic interactions describe antagonistic relationships between consumers and their resources, mutualistic interactions account for reproductive services provided to the resource species. Mutualistic interactions often describe an interaction between species where one receives a service (the service receiver) while the other receives a reward for facilitating the service (the service provider). In plant-pollinator interactions, this service is reproductive, where the pollinator delivers pollen to the stigma of female plants, while the reward is energetic, often in the form of nectar. This results in a fitness benefit to both species, and plays a fundamental role in generating and maintaining local biodiversity [17]. Mutualistic interactions can serve to minimize competition between species [18, 19] and in some cases increase the diversity of the community [20]. The abundance of mutualistic interactions can be substantial in many systems. For example, more than 90% of tropical plants depend on animals for dispersing their seeds, and many plants depends on animals to pollinate their flowers [21, 22].

Mutualistic systems can be depicted by a bipartite network [23], consisting of two sets of nodes – here denoting service providers, or pollinators, and service receivers, or plants – and where links between them identify a mutualistic dependence. These networks have been observed to display consistent patterns in their structure, suggesting a common yet enigmatic process governing their assembly [24–27]. Specifically, networks of mutualistic species tend to have a highly nested structure, where specialist species interactions tend to be subsets of generalist species interactions [28–30]. Nestedness is an important structural characteristic, as it lowers the intensity of interspecific competition, increasing the number of species that can coexist [19], promoting diversity. Of particular importance in island systems — where species arrivals and extinctions occur at an accelerated pace — is how these dynamics influence the structure and function of mutualistic communities. Understanding how such dynamics shape the nested architecture of mutualistic networks can shed light on the processes that support ecological resilience in island ecosystems.

Compared to mainland mutualistic systems, those on islands are less diverse, have a lower ratio of pollinator to plant species and a higher number of supergeneralists (*i*.*e*.species with many interactions)[31–33]. These supergeneralists can play a critical role in shaping the structure of mutualistic networks by increasing the density of interactions (*i*.*e*., higher connectance) and promoting network nestedness [32, 34, 35]. For example, Trøjelsgaard et al. [36], used island ages as a proxy for the temporal assembly process and showed that recently formed island communities hosted a larger number of generalist pollinators, whereas more established island communities had larger numbers of specialized pollinators and plants.

Remote oceanic islands have provided unique opportunities to integrate ecological and evolutionary dynamics, enhancing our understanding of how diverse ecological communities are formed and maintained [37]. On islands, speciation plays a crucial role in generating species diversity, with island communities shaped by the interplay of immigration, speciation, and extinction rates [2, 38]. Losos and Schluter [39] demonstrated that on smaller islands, speciation events are rare, such that immigration is the primary source of new species. However, on larger islands (greater than 3,000 Km^2^), speciation surpasses immigration as the dominant driver of species diversity, with the rate of speciation increasing proportionally with island area [38]. Additionally, the number of species generated through speciation only exceeds those arriving through immigration when the speciation rate is greater than half of the extinction rate [40]. Although very insightful, we still need to scale down to better understand how evolutionary dynamics shape species traits and alter the adaptive landscape, while communities are still being shaped by ecological dynamics, such as immigration and extinction, on islands.

As mutualistic species interact they alter the fitness landscape, influencing the selective forces operating across species [41]. And because species are connected in a network, the coevolutionary changes impacting species that do not interact directly with a particular pollinator or plant (the indirect effects) can often be just as influential as those that do [42]. These processes may feed back to alter the forces governing the dynamics of colonization and extinction, introducing a great deal of complexity to coevolving mutualistic island communities. For example, coevolution can favor trait-matching between species, such as the size of the nectar tube in plants and the size of the proboscis among insect pollinators [43–45]. Indeed, this is the subject of Darwin’s famous insight into the proboscis length of the then unknown pollinator of the star orchid *Angraecum sesquipedale* [46]. However, a key challenge lies in understanding how coevolution influences not only a pair of interacting species, but a diverse and ever-changing community. Coevolution can lead to trait matching between species of different guilds, even when they are components of a larger and more complex mutualistic network [47, 48]. Over time, the evolution of traits mediating mutualistic interactions can have profound impact on the eventual structure of the community [49–51]. In fact, trait-matching between plant and pollinating birds has been shown to be directly predictive of island network structure [48], while indirect interactions are fundamental to the coevolutionary dynamics of mutualistic networks, especially in nested communities [52].

Here we aim to understand how mutualistic networks assemble into island communities. We integrate colonization and extinction dynamics into a stochastic model of trait coevolution to uncover how the reciprocal nature of the selective forces driving trait-matching between interacting species influence the assembly process, and vice versa, to alter the composition and stability of evolving communities. Specifically, we aim to address three main questions. First, how do extinction rates influence the structure of mutualistic networks and the mean trait of species on islands? Second, does the structure of the mainland community (source) influence that of assembling island communities? And third, does coevolution, when governing the extinction process, play a significant role in shaping the structure of mutualistic networks of island communities? Our investigation contributes to a deeper understanding of community assembly processes and informs conservation efforts aimed at preserving ecosystem function and biodiversity in island systems.

## 2 Methods

We begin with an interaction network between two groups of species: those delivering a reproductive service while receiving trophic rewards (*e*.*g*. pollinators), and those receiving a reproductive service and providing trophic rewards (*e*.*g*. pollinating plants). This bipartite network is specified by the adjacency matrix **A**, where the rows *i* of the matrix represent plant species and the columns *j* represent animal pollinator species for a given mutualistic system. If two species engage in a mutualistic interaction, the matrix element *a*_*ij*_ = 1; and is zero otherwise. Throughout we will simulate evolutionary dynamics of species assembling into an island community. While the composition of species in the island community depends on those who have successfully colonized, their interactions with each other are assumed to be as observed in the full empirical community on the mainland. Within each system, we evaluate the dynamic structure of the island community as species colonize and go extinct, and as the mutualistic traits of species coevolve over time, where species’ traits are assumed to determine the efficacy of each pairwise mutualistic interaction. Each species’ trait is represented by a quantitative character where trait-matching between plant and pollinator is presumed to maximize fitness. In other words, in a simple interaction pair, without the confounding effects of the community, the traits of plant and pollinator species will eventually evolve to match, where, for example, the size of the pollinator’s proboscis will evolve to fit the the length of the plant’s nectar tube, and vice versa.

### 2.1 The dynamics of coevolutionary assembly

We investigate the combined effect of trait coevolution in a mutualistic system as it assembles and evolves from an established mainland community. Throughout, we assume that the traits of species in the mainland community have reached a coevolutionary steady state prior to the assembly process [cf. 53], and that the mutualistic structure of species present in the assembling community are invariant with respect to the mainland. So while the presence/absence of interactions between coexisting species are the same as those on the mainland, the existence of an interaction is contingent on the co-occurrence of both species in the assembling community, and the interaction is absent if one of the pair is not present on the island. To establish mainland communities, we used 50 empirical pollinator-plant networks from the Web of Life database (http://www.web-of-life.es/), introducing a wide range of network topologies and localities, ranging from Argentina to Canada (Table S1). We did not include empirical island networks, as we intended the network assemblage to represent mainland systems from which simulated island communities were assembled.

The mainland community represents the pool from which the island community is assembled. After finding the deterministic steady state of the mainland community following Guimarães et al. [53], we allowed a species *i* to assemble into a novel community, arriving with the mean trait value *Z*_*i*_, initially obtained from the corresponding steady state mainland value. While the mainland steady state is found deterministically, assembly of the island community follows a stochastic process, where species colonize, coevolve, and suffer extinction with probabilities that change with the richness of the community. Following the MacArthur-Wilson Species Equilibrium Model [54], colonization rates are assumed to decrease as the number of species on the island increase, whereas extinction rates - as well as coevolutionary rates increase with the number of colonized species. We next detail how colonization, coevolution, and extinction change with the state of the assembling community, and whether and to what extent they depend on species’ mutualistic traits.

#### Colonization

The assembly process is initiated by colonization, where a mutualist pair, consisting of both a plant and pollinator, is randomly selected from the mainland network to colonize the system. The colonization of subsequent species requires the presence of at least one mutualist partner in the island community. So while the mainland community defines the full suite of potential mutualistic partners for a given species, it is only capable of colonizing the island community if at least one of those potential partners is present on the island. Each colonization event consists of a single species being transported to the island system. Once the species colonizes the island, it realizes the subset of its potential mutualistic interactions allowed by its mutualistic partners in the island community, potentially facilitating additional future colonizations.

#### Coevolution

The trait coevolutionary process follows that of Guimarães et al. [53], where we track the mean value of a quantitative trait for each species 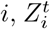, which evolves in one generation *t* to the next (*t* + 1), and is initialized from the species-specific steady state mainland value. The trait *Z*_*i*_ is assumed to have heritability *h*^2^ and its total additive phenotypic variance in the population is given by 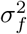, which is assumed to be the same across species. Heritability is defined as 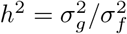, where 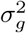 is the total additive genetic variance, and we assume throughout that 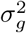 is constant over time and equal for all species. Trait values for each species change over time due to both mutualistic selection exerted by species *j*, denoted as *M*_*ij*_, and environmental selection on species *i, E*_*i*_, such that the trait dynamic can be written in discrete time as

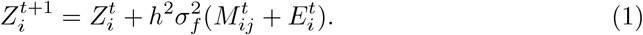

We assume that mutualistic selection favors trait-matching, such that

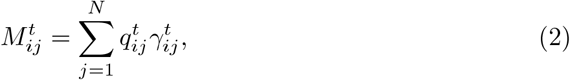

where *γ*_*ij*_ is the ‘trait mismatch’ between species *i* and *j*, where 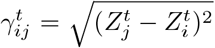.

We note that 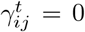 if the trait values of interacting species are equally matched, reaching a fitness maximum, and 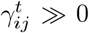 if they are mismatched. 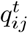 is the impact that each interaction has on the mutualistic selection differential, such that

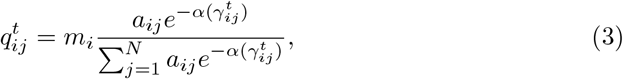

where *m*_*i*_ is the strength of mutualistic selection, *a*_*ij*_ is the corresponding element of the adjacency matrix, and 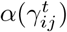 is a scaling parameter controlling the sensitivity of mutualistic selection to trait matching. In other words, *q*_*ij*_ is the relative evolutionary effect of species *j* on species *i* in relation to all of its interacting partners, ranging from 0 (no interaction) to 1. Because the island community is a subset of that of the mainland, the mutualistic selection (*M*_*ij*_) exerted on species *i* will differ accordingly, favoring trait-matching of species present in the island community at a given point in time (see Eq. 2), diverging from the mainland steady state where all species are present.

The environmental effect on the mutualistic trait for species *i* selects towards an environmental optimum *θ*_*i*_. For simplicity, we assume that species *i* on the mainland and the island will have the same *θ*_*i*_ value, suggesting that the mainland and on the island environments are similar. Environmental trait optima are randomly chosen from a uniform distribution between 0 and 1 for each species and kept constant over the course of a given simulation. The environmental effect on selection is then defined as

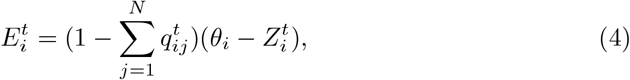

where 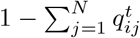 is the strength of the environmental selection.

Combining equations 1, 2, 4, along with 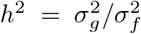 we obtain the full trait dynamic

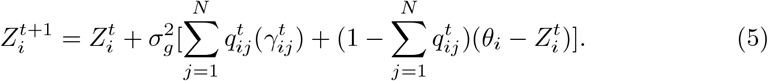

#### Primary extinction

Local extinctions are introduced as a stochastic process, where species are removed from the island community, but can recolonize at a later point in time. We distinguish primary extinctions from secondary extinctions, the latter of which occurs subsequent to a primary extinction. We examined two primary extinction scenarios: random extinctions, where all species have the same probability of going extinct, and trait-based extinctions, where the probability of extinction increases with the mismatch of plant and pollinator traits (higher *γ*_*i*_).

Following optimal foraging assumptions [55], we assume that increased mismatch between a mutualistic pair results in lower interaction efficiency. We can formalize this notion by assuming that lower interaction efficiency means a longer average handling times performing mutualistic services, reducing the species’ net reward [56–58]. While we do not simulate individual interactions in our model, we can use this formalization to derive the probability of extinction as a function of trait mismatch between species *i* and *j, γ*_*ij*_. If we assume that mutualistic interactions between a pollinator *i* and its set of mutualistic partners results in some net profitability (measured as either energetic or reproductive reward), its expectation can be written as

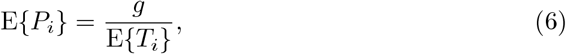

where we formalize E{·} as an expectation and denote *g* as the gain, assumed to be constant, and E{*T*_*i*_} as the expected time a species spends in a pollination interaction. The time a pollinator *i* expends visiting *N* partner species, *T*_*i*_, is a function of its trait mismatch relative to the plants *j* it is pollinating, and can be written as

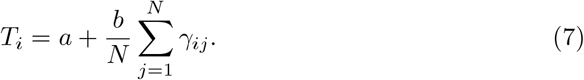

Here, *a* represents the mean handling time if mutualistic partners are perfectly matched (*γ*_*ij*_ = 0), and *b* represents the effect of dissimilarity on increases to the temporal cost, where for simplicity we assume a linear relationship. We also assume for simplicity that each potential mutualistic partner has equivalent values of *a* and *b*.

As species’ traits vary, the dissimilarity of a pollinator’s trait with respect to its mutualistic partners can be treated as a random variable. A species *i* thus has a distribution of dissimiliarity values with its set of mutualistic partners, and because these values are constrained between 0 and 1, we can assume they can be described by a beta distribution with expectation E {*γ*_*i*_ } and variance Var {*γ*_*i*_} . From this, we first derive the expected interaction time a pollinator spends with its mutualistic partners, E{*T*_*i*_}, and approximate the expected profitability as

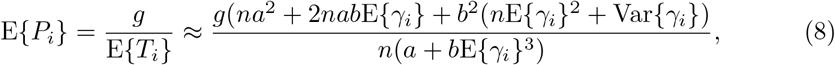

where the expectation and variance of a species’ dissimilarity with its interacting partners describes its dissimiliarty distribution at a given point in time (see Supplemental Materials for a detailed derivation). While we have illustrated derivation of the expected profitability for a mutualistic service providers, the profitability for service receivers can be estimated in the same way, though with the appropriate change in units, which may be reproductive rather than energetic. We note that this specification serves two important purposes. First, it provides a mechanistic linkage between a quantity with specific ecological and energetic importance (profitability) to the more abstract notion of trait mismatch. Second, while we do not directly connect energetic parameters to empirical systems here, Eq. 8 provides a means to do so, bridging the theory explored here to potential application to mutualistic communities in nature.

The expected profitability for species *i*, E {*P*_*i*_}, is the expectation of a profitability distribution that remains unspecified. Regardless, we can assume that as the expected profitability declines (as, for example, mutualistic partners become more mismatched), the probability of extinction will increase. If the the unspecified profitability distribution is Gaussian in nature (where 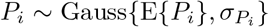), the probability that a species’ profitability will fall below some critical threshold *χ*, inducing extinction, can be written

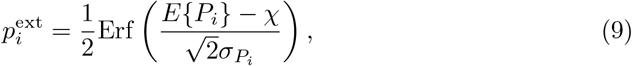

where Erf() is the error function. This exercise allows us to directly translate the dissimilarity distribution for a species *i* to a probability of extinction as a function of trait mismatch, which in this case increases sigmoidally with trait dissimilarity.

#### Secondary extinction

Following primary extinctions, which could either be determined by randomly selecting species for extinction, or by calculating the probability of extinction based on trait mismatch (Eq. 9), secondary extinctions may result. Secondary extinction occurs when a species loses all of its interacting partners, leaving it disconnected from the island community. In the case of a pollinator, this means that its energetic gain is eliminated; in the case of a pollinating plant, this means that its reproductive potential cannot be met. We assume that secondary extinctions follow primary extinctions in the same time step, implicitly assuming plants are annual, rather than perennial species. Moreover, this means that mutualistic interactions are facultative, except in the case where only a single interaction remains. By including both primary and secondary extinctions, we acknowledge the possibility of extinction cascades, where the loss of a single species can ripple through and cause extinctions across many others.

### 2.2 Stochastic assembly algorithm

We implement colonization, coevolution, and extinction processes using a Gillespie Algorithm (see Supplementary Materials for details), where the rates *r*_*j*_*q*_*j*_ of all possible events *j* (here, colonization, coevolution, and extinction) are computed in a given step, where *r*_*j*_ are assigned constants to modify the likelihood of event *j*, and where *q*_*j*_ represents the number of species prone to event *j*. The time at which the next event happens is calculated by drawing a random number from an exponential distribution with mean 1*/* ∑_*j*_ *r*_*j*_*q*_*j*_. A randomly selected event then occurs from the set of possible events such that the probability of event *x* is *q*_*x*_*r*_*x*_*/* ∑_*j*_ *q*_*j*_*r*_*j*_, whereas the time interval over which this occurs is given by Δ*t* = 1*/*∑ *j q*_*j*_*r*_*j*_. The effect of the event is then implemented and the list of possible events is updated for the next step. This algorithm offers a much better approximation to the true stochastic continuous time process than a simulation in discrete time steps, while providing a much higher numerical efficiency [59].

To explore the dynamics of the system, we simulated each network for 5000 time steps (where an event occurs at each step in accordance with the Gillespie algorithm), which we visually confirmed was enough to avoid transient effects. For each of the 50 empirical mainland plant-pollinator networks, we implemented 1000 replicates for both random and trait-based extinction scenarios and our results report the mean of these 1000 replicates for each network. Throughout, we set the rate of colonization *r*_c_ = 1 and the rate of evolution *r*_evol_ = 1. We varied the effect of increasing primary extinction rates relative to those set for colonization and coevolution, from low (*r*_ext_ = 0.3), to medium (*r*_ext_ = 0.6), to high (*r*_ext_ = 0.9). For the summary of model parameters see Table 1.

**Table 1:**
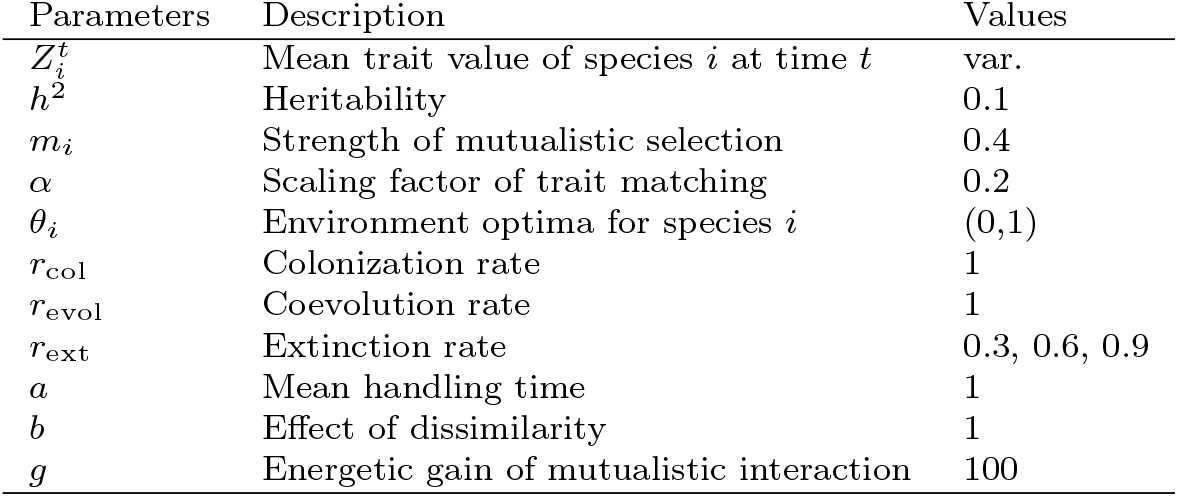
Parameters, descriptions, and set values or range.

| Parameters | Description | Values |
| --- | --- | --- |
| $Z_i^t$ | Mean trait value of species $i$ at time $t$ | var. |
| $h^2$ | Heritability | 0.1 |
| $m_i$ | Strength of mutualistic selection | 0.4 |
| $\alpha$ | Scaling factor of trait matching | 0.2 |
| $\theta_i$ | Environment optima for species $i$ | (0,1) |
| $r_{\text{col}}$ | Colonization rate | 1 |
| $r_{\text{evol}}$ | Coevolution rate | 1 |
| $r_{\text{ext}}$ | Extinction rate | 0.3, 0.6, 0.9 |
| $a$ | Mean handling time | 1 |
| $b$ | Effect of dissimilarity | 1 |
| $g$ | Energetic gain of mutualistic interaction | 100 |

## 3 Results

### 3.1 The structure of island communities

Our theoretical framework generally reveals fast initial assembly of the community, where species richness *S* sharply increases to oscillate around a steady state value *S*^*^, the value of which varies across mainland networks (Figure 1A and S1). Because different mainland communities range widely in species richness, we compare the effects of assembly across networks by assessing relative richness *S/P*, or the richness attained on the island *S* relative to that of the mainland *P* . While island communities assemble to, and oscillate around, a steady state in species richness relatively quickly, the amplitude of fluctuations also varies from network to network. While temporal fluctuations (*i*.*e. oscillations*) in relative island community richness may be expected to be larger for less diverse networks, the coefficient of variation (*CV*_*S*_), defined as *CV*_*S*_ = SD{*S*}*/S*^*^ presents a more comparable depiction of relative fluctuation size.

**Fig. 1:**
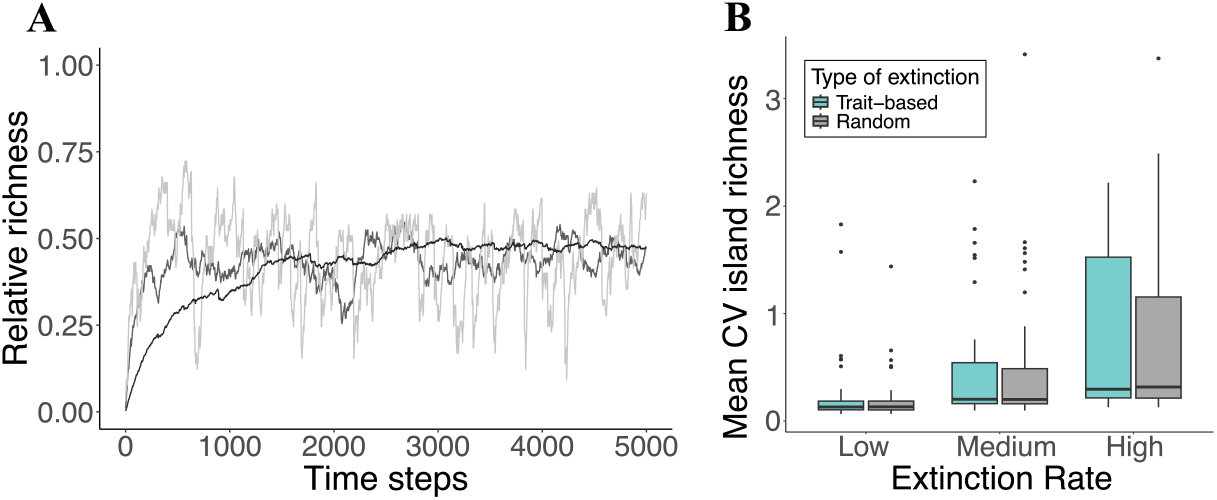
**A)** Example of island species richness dynamics over time for three different networks under medium extinction rate. Richness values of each network is relative to the source (mainland) networks **B)** A boxplot depicting the distribution of the coefficient of variation (CV) of species richness across networks, where each data point in the distribution represents the mean CV calculated from 1000 simulations, evaluated assuming low, medium, and high extinction rates

We assessed the *CV*_*S*_ for each assembled network, discarding the first 3000 event steps to avoid the effects of initial conditions. A higher *CV*_*S*_ implies greater community instability (larger fluctuations relative to the mean richness), whereas a lower *CV*_*S*_ indicates a more stable community. Our results show that a higher extinction rate leads to more unstable communities, as expected, for both the random and trait-based extinction scenario (Figure 1B and S2). For each extinction rate, we conducted Mann-Whitney U-test to determine whether there were significant differences in the *CV*_*S*_ values between the two extinction scenarios. Although both extinction scenarios have an elevated *CV*_*S*_ with increased extinction rates, we found no significant differences between the two conditions (Low: *W* = 1222, *p* = 0.88; Medium: *W* = 1137, *p* = 0.80; High: *W* = 818, *p* = 0.86). For example, when extinction rates are high, the mean *CV*_*S*_ for the trait-based extinction scenario is 0.74, whereas that of the random scenario is 0.73. Although the effects of the two different extinction scenarios are roughly similar on average, we observe slightly more variability for the trait-based extinction scenario, particularly when extinction rates are high (Figure 1B and S3).

Since the source that nourishes the island community is the mainland community, we next explore whether fluctuations in island community richness, measured as *CV*_*S*_, correlate with the structure of different mainland networks. We characterized the structure of both island and mainland communities by calculating three common measures of network structure: modularity, nestedness (NODF), and connectance [60, 61]. We used the function *networklevel* in bipartite package in R [62]. Connectance is defined by the relative link density (*L/S*^2^, where *L* is the realized number of links in the island system and *S* the number of species), whereas nestedness and modularity capture larger-scale patterns of organization in the system, and are often anti-correlated with each other. Nestedness measures to what extent more specialized species are subsets of more generalized species, where a high value indicates that such a pattern predominates. Modularity captures to what extent smaller groups of interacting species are more tightly connected to each other than to other groups, providing insight into how compartmentalized the community is. Typically, nested communities have lower modularity, and modular communities have lower nestedness. Our results show that the effect of mainland network structure on island stability, measured as *CV*_*S*_, is generally weak but is the strongest at intermediate extinction rates (Figure 2). Higher values of NODF and connectance result in greater fluctuations on the islands, while modularity has a negative relationship only at medium extinction rates (*F*_*low*_ = 2.95, *p*_*low*_ = 0.08; *F*_*med*_ = 63.12, *p*_*med*_ *<* 0.05; *F*_*high*_ = 13.82, *p*_*high*_ = 0.41). For NODF, a significant relationship is observed across all extinction rates, with the strongest effect at medium extinction (*F*_*low*_ = 10.33, *F*_*med*_ = 38.52, *F*_*high*_ = 8.17, *p <* 0.05). Similarly, connectance significantly influences stability at all rates, with the strongest correlation at medium extinction (*F*_*low*_ = 36.04, *F*_*med*_ = 63.12, *F*_*high*_ = 13.82, *p <* 0.05).

**Fig. 2:**
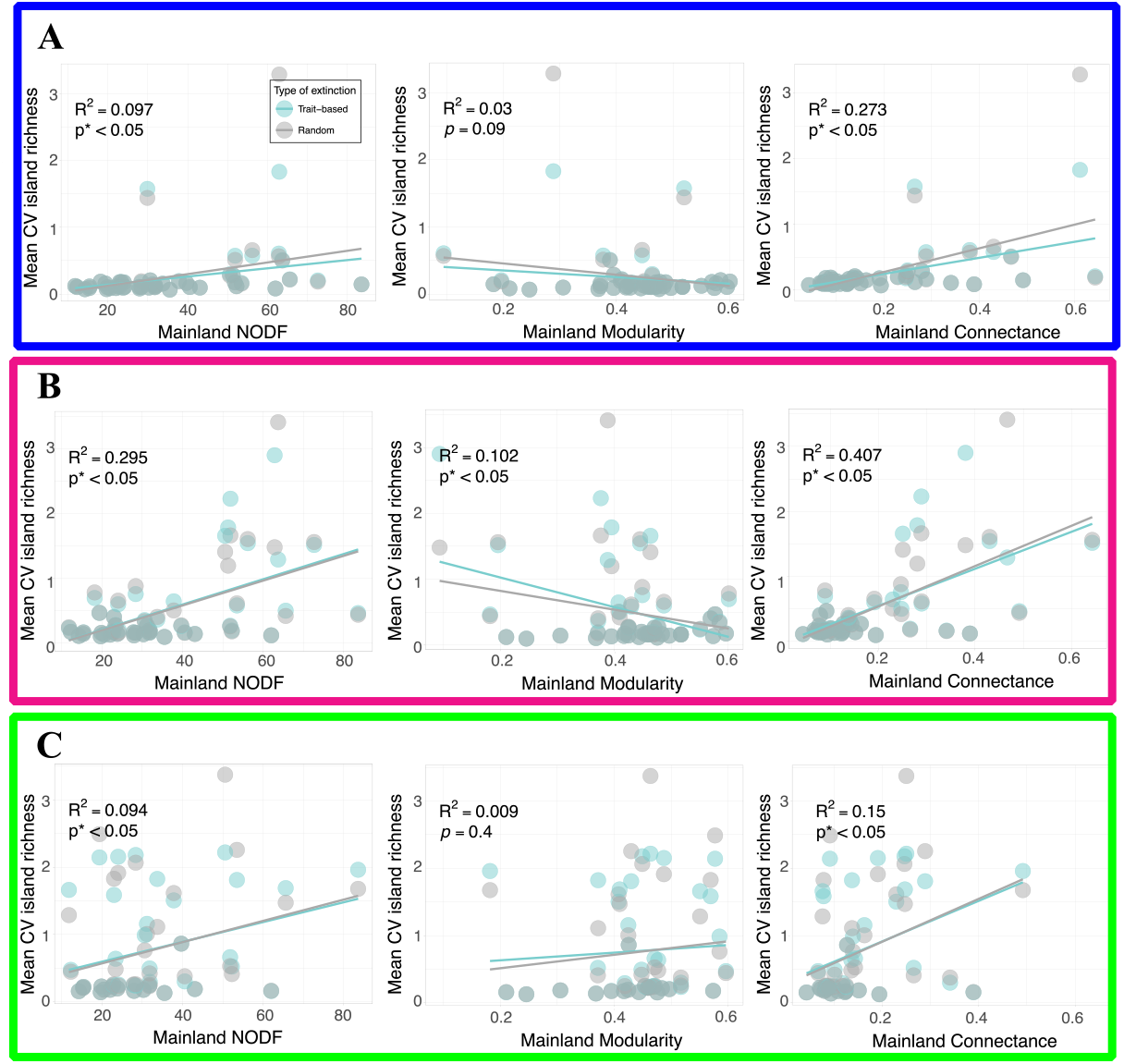
Exploring how mainland network structure affects island community instability (*CV*_*S*_) at three different extinction rates: **A)** Low (blue), **B)** Medium (pink), and **C)** High (green).

As extinctions rates transition from medium to high, many communities experience complete collapse. We counted how often island communities crashed (*i*.*e*.all species went extinct) across 1,000 simulations for each network. As expected, increasing extinction rates led to more collapses in the system (Figure S4). We then investigated whether the structural features of mainland community networks measured above correlated with a higher likelihood of collapse for the island community. Our results show that the structure of the mainland does not play a great role in the number of island collapses for either extinction scenarios (Figure S5).

We next assess how the dynamic structure of islands corresponds to their dynamics. Island network structure was assessed for the last 1000 time steps of each simulation, at intervals of 100 time steps, and averaged across 1000 replicates, which we repeated for each network. The assembling island communities assemble and evolve structures that diverge from their mainland progenitors, with a relative species richness that declines with increasing extinction rate (Figure 3A). We also observe that island communities have lower modularity and a correspondingly elevated nestedness and connectedness than mainland networks (Figure 3B-D). The degree of nestedness and connectance increases with higher extinction rates, while our metric of modularity appears not to vary much across extinction rates. It is also clear that at higher extinction rates, island communities reveal lower variability in modularity (*SD*_main_ = 0.11; *SD*_isle_ = 0.07 and 0.07, trait-based and random extinction, respectively), nestedness (*SD*_main_ = 17.85; *SD*_isle_ = 11.06 and 11.12, trait-based and random extinction, respectively) and connectance (*SD*_main_ = 0.14; *SD*_isle_ = 0.09 and 0.09, trait-based and random extinction, respectively) than their mainland progenitors (see Figure 3 B-D). We performed a Kruskal-Wallis test to test the difference between the three groups, fallowed by a pairwise comparisons. We found significant difference between mainland and island structure, but the different extinction scenarios do not reveal large structural differences (Table S2).

**Fig. 3:**
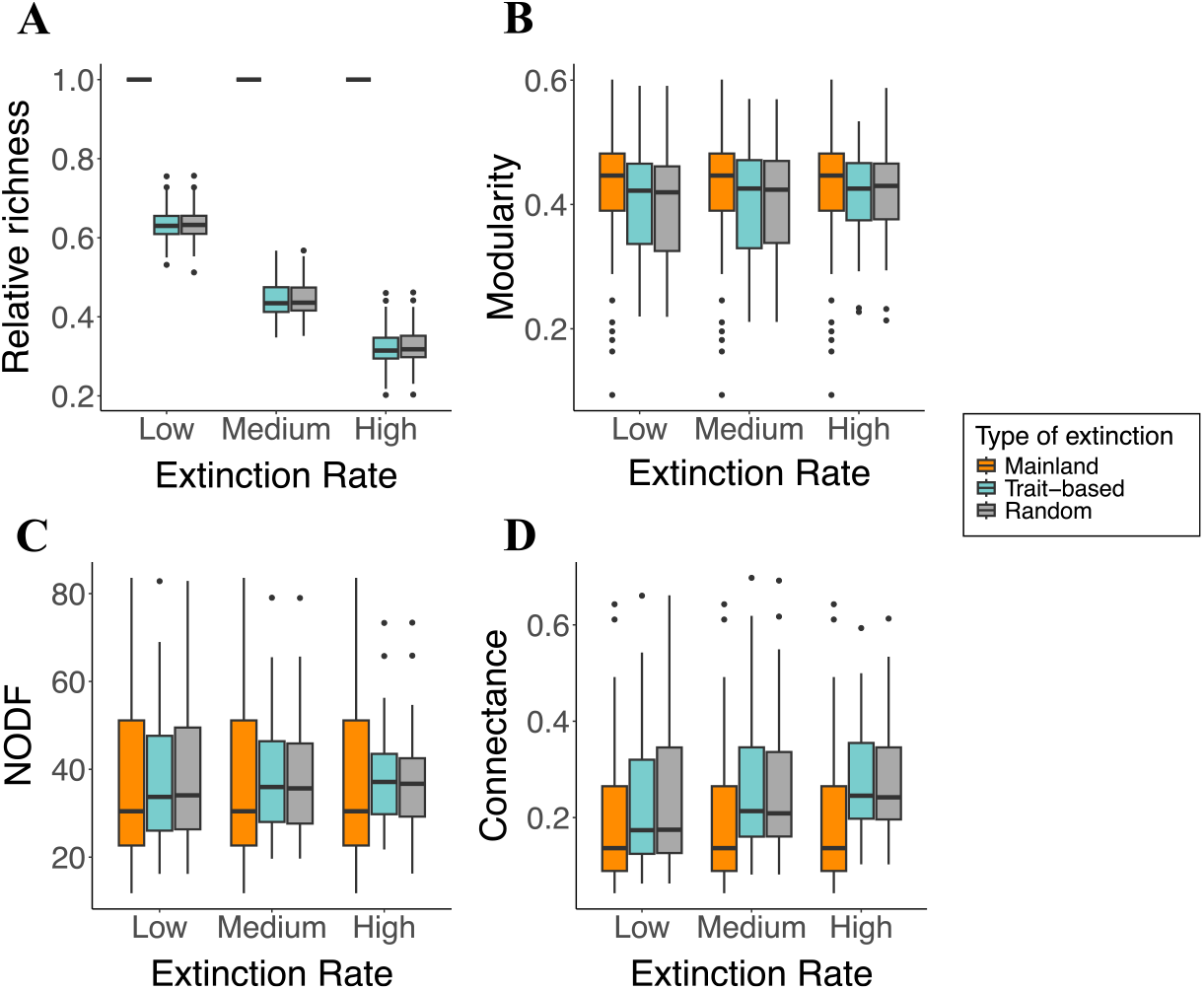
Structure of assembled networks on the island compared to the mainland for different extinction rates. Each point represents the mean from 1,000 simulations for each of the fifty pollination networks. **A)** Relative richness; **B)** Modularity; **C)** Nestedness (NODF); **D)** Connectance.

### 3.2 Species Persistence on islands

The assessment of mainland (source) and island (realized) structure illustrates how larger-scale patterns of interactions may influence the dynamics of the island community. As a next step, we assessed the role of species-specific characteristics, such as the number of interacting mutualistic partners, on these dynamics.

Since community trajectories fluctuate around different steady states *S*^*^, we calculate persistence as the percentage of each species’ occurrence on the island over the temporal span of the simulation. We examine the persistence of species within each mutualistic community relative to the number of its potential mutualistic partners in the mainland community, which we denote as the ‘potential degree’. The potential degree of each species describes its inherent ability to specialize or generalize across mutualistic partners, which may or may not be realized as colonizations and extinctions change the composition of species in island communities. We observe that a species’ persistence on the island increases sharply with its potential degree, saturating to a fixed persistence value for species with very high potential degree (Figure 4A and S6). This pattern indicates that specialist species with low degree have low persistence, as their few interactions limit their ability to weather changes in community composition across the assembly dynamic. Vitally, these low-degree species are prone to both primary and secondary extinctions, as they are more frequently losing their few partners. The steep increase in persistence with each additional interaction means that the observed ‘generalization advantage’ saturates at a relatively low interaction degree.

**Fig. 4:**
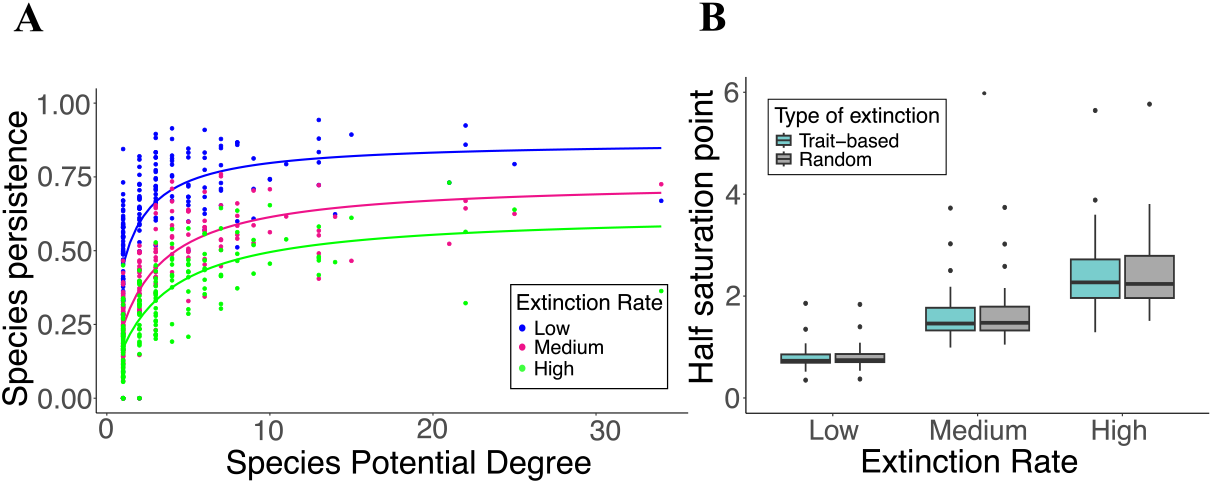
**A)** Species persistence relative to the potential species degree for a single island community under low, medium, and high extinctions rates. Each point is a species and we used the same mainland network as the source. **B)** A boxplot depicting the distribution of the approximated half saturation across networks, where each data point in the distribution represents the mean calculated from 1000 simulations, evaluated assuming low, medium, and high extinction rates.

We can determine the number of interactions corresponding to 50% of the maximum persistence across low, medium, and high extinction rates by identifying the half-saturation point of a saturating function fitted to the persistence data (lines in Figure 4A). We observe the mean of the approximated half saturation points in units of degree is ca. 0.8 assuming a low extinction rate, with effectively no difference between trait-based and random extinction scenarios (Fig. 4B). A low half-saturation degree means that a relatively small number of interactions promote a substantial increase in persistence. In this case, because the half-saturation degree is *<* 1, a single interaction enables, on average, persistence in the system for *>* 50% of the evaluated simulation time. As the rate of extinction increases, the potential for even generalists to persist is eroded, and the half-saturation degree increases to ca. 2.51 (Fig. 4B). Accordingly, increasing extinction rates increases the number of interactions needed to maximize species’ persistence on islands. Lastly, our results indicate that as extinction rates increase, the variability in the half-saturation degree also increases, demonstrating that certain networks require more interactions than others to reach the 50% threshold of maximum persistence.

### 3.3 Coevolutionary assembly dynamics of island communities

To better understand the effect of coevolution on assembled island communities, we calculated average trait dissimilarity taken across the community 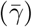. We computed the average species trait dissimilarity 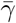 at the last time step of our simulation across 1000 replicates for all mainland networks under both extinction scenarios, and across increasing extinction rates. At the community level, a low mean trait dissimilarity value means that species on the island community have traits that are, on average, more similar. While the random extinction scenario removes species randomly, regardless of trait dissimilarity, the trait-based extinction scenario assumes species more dissimilar than their mutualistic partners are more prone to primary extinctions.

First, we observe that the interplay between coevolutionary dynamics and the assembly process results in island communities that have average trait dissimilarity values that differ greatly from their mainland counterparts, and this difference grows as the extinction rates increase (Figure 5A). Second, there is a clear difference in 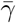 between trait-based and random extinction scenarios. The ANOVA test showed that mainland, island with trait-based, and random extinction differed significantly across all three extinction rates (*F*_*low*_ = 5.31, *F*_*med*_ = 1.82, *F*_*high*_ = 9.51, *p <* 0.05)and the post-hoc test further revealed significant differences between the three groups. As the extinction rate increases, the mean dissimiliarty of mainland and island communities under the random extinction scenario also increases (Figure 5A). When extinctions are a function of trait dissimilarity, we observe that the mean dissimilarity values are much lower in island systems compared to mainland communities (Figure 5A). Over the course of the assembly process, species that are dissimilar are more likely to suffer extinction, with communities becoming increasingly similar. Because both the trait-based and random extinction scenarios employ the same coevolutionary process, which favors trait matching, the decrease in dissimilarity in trait-based extinction communities can be interpreted as the result of the assembly, rather than the coevolutionary dynamic.

**Fig. 5:**
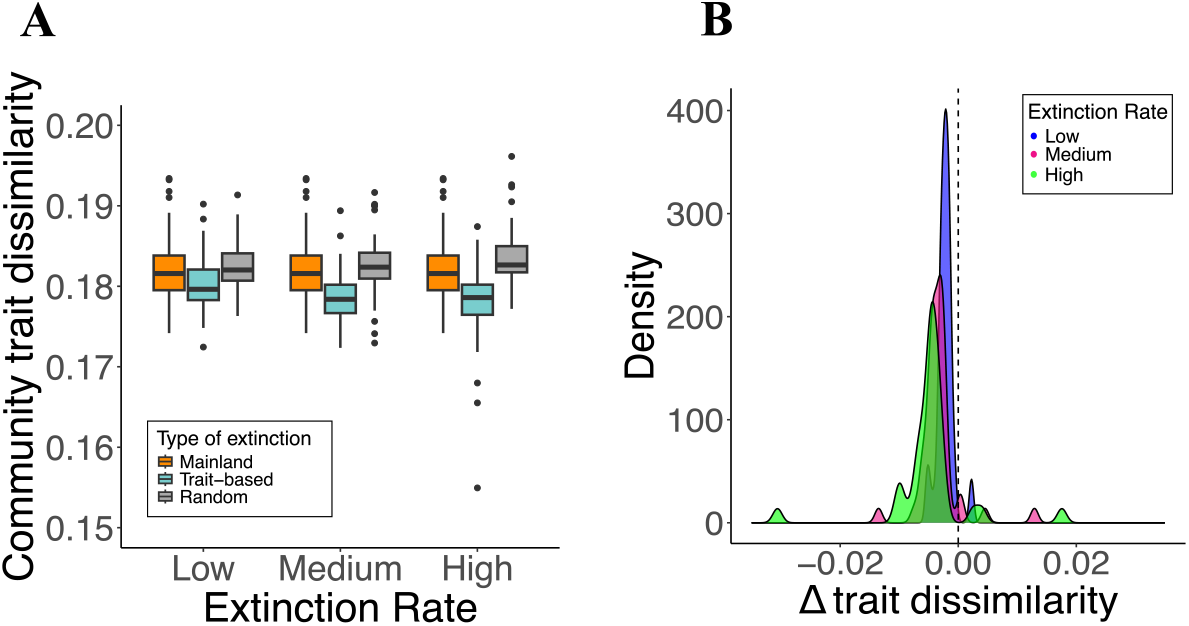
**A)**Higher extinction rates decreases trait mismatching among species on the island community. Trait dissimilarity is the mean trait mismatch of the island community of 1000 simulations across 50 networks. **B)** Δ is the paired difference of trait dissimilarity of the same networks in a trait-based minus the random extinction scenario for the 50 pollination networks. Low, medium, and high extinction rates represent *r*_ext_ = (0.3, 0.6, 0.9), respectively.

Second, we note that the assessment of community dissimilarity across networks may mask to what extent individual networks increase or decrease in dissimilarity as extinctions move from random to trait-based. Comparing the difference between community dissimilarity values for the random and trait-based extinction scenarios on a per-network basis, where 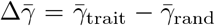, negative values indicate that networks assembling under the trait-based extinction scenario result in lower community dissimilarity, while positive 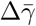indicates that the trait-based extinction scenario results in higher community dissimilarity values. We observe a clear shift to lower negative values for 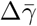 from low to high extinction rates (Fig. 5B). This result demonstrates that on a per-community basis, the effect of trait-based extinctions serves to lower the trait dissimilarity of interacting species, an effect that grows with increasing extinction rates.

## 4 Discussion

We investigate the assembly and coevolution of island communities from a mainland source pool by integrating a coevolutionary model premised on trait-matching with a stochastic assembly dynamic. By combining these approaches we aim to disentangle the potential influence of both ecological and evolutionary dynamics in shaping the structure and function of assembling island communities.

Beyond capturing the expectation that island richness is inversely related to the rate of extinction, we also observed that the relative size of fluctuations in richness (measured as *CV*_*S*_) over the course of assembly increased with extinction rate, while differing widely among island communities (Figure 1 and S3). When the extinction rate is low, the island can assemble a larger range of species, however the assembly dynamics and resulting fluctuations may be relatively insensitive to both structure (interactions) and function (trait matching between mutualistic partners). As the extinction rate increases, the community reaches a steady state composed of fewer species, though the dynamics may increasingly reflect the selective process being imposed. When extinction rates are too high, the island is in a constant state of flux – from both the primary extinctions but perhaps even more-so the secondary extinction cascades. Higher extinction rates also led to networks that were more nested and connected, but less modular, providing insight into the roles of structure in determining dynamics. The idea that the structural properties of ecological networks — such as connectance, modularity, and nestedness — play a crucial role in determining the robustness of ecological communities, *i*.*e*. how the community respond to a extinction cascade affect, have been explored for different type of ecological interactions, with mixed results. For instance, connectance has been shown to either increase or decrease robustness depending on specific model assumptions [63–65], while modularity can enhance robustness by confining cascading effects to specific modules or, conversely, increase vulnerability within those modules [66]. Nestedness, however, is often considered a proxy for robustness [67], though the positive relationship between nestedness and robustness has only been clearly demonstrated when specialists have a higher probability of extinction than generalists [68]. The observation that increased extinction rates contribute to a wide range of *CV*_*S*_ across networks suggests that inherent differences in network structure play a key role in generating destabilizing fluctuations. Notably, when species extinctions are influenced by trait dissimilarity, this variability in the relative size of fluctuations is amplified compared to when extinctions are random (Figure 1B). This indicates that while coevolutionary processes within our framework help reduce trait dissimilarity, extinction events based on trait differences can heighten fluctuations in certain island communities, resulting in less stable systems.

Because island systems pull from the species and associated structures of mainland communities, the notion that multiple spatial scales contribute to the regional processes shaping community assembly [69–71] is inherently captured within our assembly framework. Here, the structure of the mainland communities giving rise to assembled islands provides insight into the potential interactions available to species, the realization of which depends on the state of the island community at a given point in time. Our result shows that when extinction rates are intermediate – not so low that they have little effect, and not so high the community is constantly crashing – the influence of mainland structure on island communities is strongest. Mainland communities that are more nested, more connected, and less modular tend to generate island communities that are more unstable (Figure 2B). That both increased connectance and nestedness correlate with larger fluctuations, while increased modularity has the opposite effect, and that there is effectively no difference between trait-based and random extinction scenarios, suggests three potentially intertwined effects. First, mainland structure does not appear to determine observable differences between systems with and without trait-dependent extinctions, suggesting the relationships are instead primarily driven by the assembly process rather than coevolution-mediated changes in traits. Second, small-world type interaction structures, where the indirect paths connecting species are short, may more effectively drive larger extinction cascades, as the influence of one species’ extinction can directly impact more species in the network [72, 73]. Third, and the converse of the second effect, such cascades are likely dampened by more modular networks, where the impact of a cascade may be severe within a module, but be less likely to expand outside of it [74, 75].

We also demonstrate that the persistence of supergeneralists is not incredibly higher than that of generalists with fewer interactions, and this is reflected by the saturating function that best fits persistence as a function of species’ degree (Figure 4A). Specialists, on the hand, have a higher turnover rate. Because specialists can more readily evolve to match their partner’s trait value, this suggests that it is the ecological processes – a combination of both the ability to colonize and a resistance to secondary extinction cascades – that instead shapes the generalist advantage. Moreover, our model prediction that benefits saturate with increasing numbers of interactions is a pattern that is also observed in natural systems. Olesen et al. [76] explored the temporal dynamics of mutualistic networks over a span of 12 years and uncovered a similar pattern: species with fewer interactions had higher turnover rates, and while those with more interactions persisted longer, the gains followed a saturating relationship, such that larger numbers of interactions delivered diminishing returns. A common assembly process proposed by Barabási and Albert [77], known as “preferential attachment”, describes the assembly of nodes in a network that tend to attach to those with many connections, where the ‘rich get richer’. Processes that are analogous to preferential attachment can also govern network disassembly, where less-connected species are more prone to local extinction, and this has been shown to give rise to more nested networks [78]. While our assembly process does not follow a preferential attachment algorithm as potential interactions are predetermined by a mainland source, less connected species are more prone to secondary extinction cascades, potentially contributing to the increased nestedness we observed in our assembling island communities (Figure 3). Intriguingly, this tendency for less-connected species to be lost during network disassembly has also been observed in empirical mutualistic networks [79, 80].

Our work advances upon previous findings by integrating the ecological processes of colonization and extinction with the dynamics of coevolution into a unified framework, where the emergence of island structure and dynamics from a mainland progenitor can be evaluated with respect to the influence of both. Empirical observations of mainland versus oceanic island mutualistic systems have not revealed clear structural differences between them, as oceanic islands are less diverse, but do not show significant differences in the connectance, nestedness, or modularity of their interactions compared to mainland systems. [32]. Importantly, they compared many island networks to mainland networks, without taking into account pairwise comparisons (mainland/source–island) or characteristics such as island size and distance from the mainland, both of which are known to impact the balance between colonization and extinction [1]. Along these lines, our framework shows that structural differences between mainland and island systems can be sensitive to extinction rate, where structural divergence between mainland and island systems increase with the rate of extinctions (Figure 3). When extinction rates are low, the similarity between island and mainland community structure increases (Figure 3B-D).

Lastly, our results show that coevolution leads to greater trait-similarity in island communities experiencing trait-based extinctions, particularly when extinction rates are high (Figure 5). Despite high species turnover, we observe that the coevolutionary dynamic serves to promote the persistence of species with similar traits, resulting in communities with greater trait-matching. However we do not find this trait-matching dynamic to leave a clear imprint on the assembled structure of island communities, as is demonstrated by the similarity of assembled islands with random versus trait-based extinctions (Figure 3). We note that coevolutionary effects in our framework may be lessened by the fact that each simulation consists of only a single mainland source and island community. As such, following a given species extinction, its recolonization effectively resets its trait value to that of the mainland, erasing any memory of previous evolutionary change. In natural systems where the island is part of a larger archipelago, there may be a greater likelihood of an ecological rescue by other island populations, connected by dispersal, that have retained this coevolutionary memory. By preserving the memory of coevolutionary processes, such dynamics could magnify the effects of coevolution and its influence in the assembly process, and this would be an obvious next step for future work.

Although studies directly linking coevolution to community assembly are limited [but see 81], there have been substantial efforts demonstrating how coevolution operates across varying spatial scales. Because local communities experience different selection pressures, coevolutionary dynamics at larger spatial scales can generate a geographic mosaic [82–85]. Geographic mosaics can function to maintain a larger diversity of phenotypes in a given region, directly impacting how the structure of mutualistic interactions varies across time and space [85–87]. Because spatial mosaics are connected via dispersal, each serves to provide a flow of new species or phenotypes to the other. For example, Medeiros et al. [88] used a mathematical model to show that increasing gene flow between mutualistic communities leads to higher trait-matching and trait convergence between interacting species. In other cases, when two populations evolve in response to very different environmental optima, gene flow can serve to flood populations with suboptimal phenotypes, resulting in migrational meltdown [89, 90]

The traits of plants and pollinators involved in pollen delivery and sequestration directly impact both the time and energy invested in each interaction, such that lower dissimilarity promotes pollination efficiency [91], and this is expected to increase fitness [87]. That island networks with random extinctions are more dissimilar, and those with trait-based extinctions are less, clearly demonstrates that coevolutionary forces are shaping the nature of species interactions (Figure 5B). Yet, as we have stated, this coevolutionary process does not leave a clear signature in the structure of island communities. Because only primary extinctions are a function of trait dissimilarity between interacting partners, we suggest that it is not primary extinctions but secondary extinction cascades that drive the structural differences we observe between islands and their mainland source communities (Figure 3B-D). So while coevolutionary dynamics shape the trait landscape within the community, on average pushing the system towards trait-matching, whatever structural imprint such a process might leave is erased by the large footprint of cascading extinctions following an initial extinction event.

By combining both the ecological processes of assembly with coevolution, we show that island communities, characterized by smaller size, and higher instability and turnover, can provide important insights into the drivers of structure and dynamics. Importantly, we show that the effects of coevolutionary trait-matching in complex mutualistic systems can be subtle, yet persistent, when communities are undergoing continuous reorganization and shuffling. Island communities contain an important source of biodiversity, and understanding the interplay between the ecological and evolutionary processes at work is crucial for elucidating the factors driving species diversity, assembly, and dynamics in relatively isolated habitats. This knowledge not only enhances our comprehension of biodiversity patterns but also informs conservation efforts by providing insights into how to preserve and manage these unique ecosystems in the face of environmental challenges and human impacts.

## S1 Supplementary information

### S1.1 Analytical approximation

We performed an analytical approximation to better understand how trait mismatch related with the probability of extinction (*p*). In our model, *t* is the time a species *j* spend interacting with species *i*. We define *t* as:

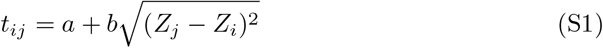

where *a* is the time if perfectly match and *b* is the effect of dissimilarity on time.

We assume these are the same for all species.

We can then calculate *T*, which is the total time a species spend in a mutualistic interactions with other species.

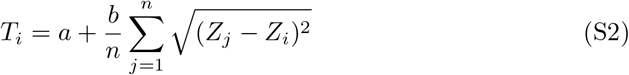

where *n* is the total number of species who species *i* interacts with.

Let define 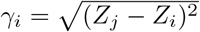. We will treat *γ*_*i*_ as a random variable and it will be drawn from a *Beta Distribution* with shape parameter *α* and *β*.

From the *Beta Distribution* we have:

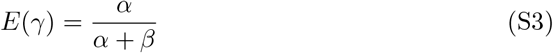

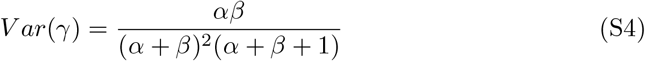

From S3 and S4 we can calculate the *E*(*T*_*i*_):

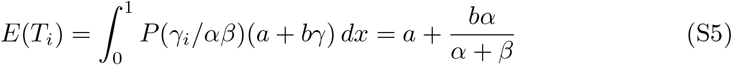

Because *V ar*(*T*) = *E*(*T* ^2^) − *E*(*T*)^2^, we then find:

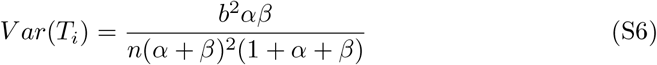

Now we define our profitability (*P*) as 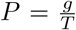, where *g* is the energetic gain of a mutualistic interaction and we assume to be constant among species. Thus, to calculate the expected profitability (E(P)) we we use the *Delta Method* approximation. Where:

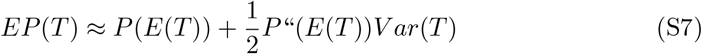

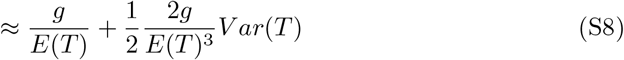

Combining S5 and S6 in S8, we have:

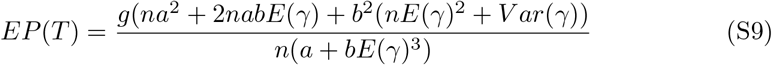

### S1.2 Tables

**Table S1:** Empirical networks used as mainland sources.

| Network | Richness | NODF | Modularity | Connectance |
| --- | --- | --- | --- | --- |
| 1 | 184 | 14.23873549 | 0.475424383 | 0.042944053 |
| 2 | 64 | 37.81390987 | 0.407417048 | 0.229166667 |
| 3 | 204 | 22.58099947 | 0.458793211 | 0.090114943 |
| 4 | 103 | 43.05299921 | 0.305314438 | 0.151371308 |
| 5 | 143 | 19.68300426 | 0.367755818 | 0.093370445 |
| 6 | 124 | 20.49405685 | 0.573327974 | 0.080362727 |
| 7 | 65 | 18.16607981 | 0.600920447 | 0.088636364 |
| 8 | 94 | 22.97842332 | 0.569003642 | 0.078282828 |
| 9 | 56 | 52.24284773 | 0.371166334 | 0.265151515 |
| 10 | 80 | 11.78630001 | 0.55047619 | 0.076252723 |
| 11 | 106 | 15.29377094 | 0.496383958 | 0.072544643 |
| 12 | 96 | 12.26008706 | 0.595800304 | 0.066869301 |
| 13 | 39 | 51.31313131 | 0.395124717 | 0.28 |
| 14 | 46 | 40.53764617 | 0.516776554 | 0.340686275 |
| 15 | 49 | 24.01294151 | 0.486968144 | 0.190883191 |
| 16 | 49 | 28.3718074 | 0.449011446 | 0.246031746 |
| 17 | 109 | 22.00459636 | 0.394785981 | 0.10791038 |
| 18 | 59 | 62.00698363 | 0.210409359 | 0.389147287 |
| 19 | 204 | 30.29379718 | 0.417036302 | 0.124039133 |
| 20 | 60 | 19.36751796 | 0.57671875 | 0.091428571 |
| 21 | 103 | 23.38299422 | 0.476938056 | 0.135817308 |
| 22 | 53 | 31.15384615 | 0.424567605 | 0.163003663 |
| 23 | 112 | 27.94153197 | 0.443159279 | 0.122660295 |
| 24 | 25 | 83.58751609 | 0.181923222 | 0.491666667 |
| 25 | 63 | 65.61027603 | 0.409322792 | 0.24691358 |
| 26 | 21 | 51.88679245 | 0.37633218 | 0.288461538 |
| 27 | 25 | 56.18849206 | 0.445331484 | 0.43 |
| 28 | 35 | 63.72412774 | 0.388352222 | 0.466666667 |
| 29 | 35 | 62.86346612 | 0.093526577 | 0.38 |
| 30 | 113 | 30.59627041 | 0.584480227 | 0.139146568 |
| 31 | 39 | 53.49656713 | 0.429576574 | 0.288888889 |
| 32 | 23 | 29.88410596 | 0.519705718 | 0.264705882 |
| 33 | 13 | 11.9047619 | 0.163518008 | 0.388888889 |
| 34 | 10 | 62.96296296 | 0.288101962 | 0.611111111 |
| 35 | 15 | 72.65306122 | 0.195834711 | 0.642857143 |
| 36 | 51 | 50.56889673 | 0.463412698 | 0.25 |
| 37 | 105 | 19.8495749 | 0.459036859 | 0.070306363 |
| 38 | 95 | 22.84928435 | 0.470859341 | 0.08234127 |
| 39 | 116 | 23.80843113 | 0.424446998 | 0.081774082 |
| 40 | 93 | 24.14551257 | 0.46889382 | 0.098484848 |

**Table S1:** (continued)
| Network | Richness | NODF | Modularity | Connectance |
| --- | --- | --- | --- | --- |
| 41 | 83 | 39.67637728 | 0.425227076 | 0.126865672 |
| 42 | 93 | 27.86605248 | 0.453690007 | 0.085714286 |
| 43 | 81 | 31.85541898 | 0.418353686 | 0.117691154 |
| 44 | 93 | 31.74693878 | 0.461230487 | 0.088900862 |
| 45 | 75 | 31.82727497 | 0.447642911 | 0.101080247 |
| 46 | 71 | 28.37614854 | 0.516888722 | 0.124794745 |
| 47 | 77 | 51.81188441 | 0.470538313 | 0.143442623 |
| 48 | 51 | 33.71946622 | 0.371613243 | 0.137254902 |
| 49 | 141 | 15.39290372 | 0.483599322 | 0.087693441 |
| 50 | 106 | 35.51303715 | 0.245710571 | 0.193859649 |

**Table S2:** Pairwise comparisons of NODF, modularity, and connectance values among mainland, trait-based (TB), and random (R) extinction scenarios using the Wilcoxon rank sum test with Bonferroni correction. Significant p-values (*<* 0.05) are in bold.

| Metric | Category | Comparison | p-value |
| --- | --- | --- | --- |
| NODF | Low | M-TB | **<0.05** |
|  |  | M-R | **<0.05** |
|  |  | TB-R | 1 |
|  | Medium | M-TB | **<0.05** |
|  |  | M-R | **<0.05** |
|  |  | TB-R | 1 |
|  | High | M-TB | **<0.05** |
|  |  | M-R | **<0.05** |
|  |  | TB-R | 1 |
| MOD | Low | M-TB | **<0.05** |
|  |  | M-R | **<0.05** |
|  |  | TB-R | 1 |
|  | Medium | M-TB | **<0.05** |
|  |  | M-R | **<0.05** |
|  |  | TB-R | 1 |
|  | High | M-TB | **<0.05** |
|  |  | M-R | **<0.05** |
|  |  | TB-R | 1 |
| Connectance | Low | M-TB | **<0.05** |
|  |  | M-R | **0.05** |
|  |  | TB-R | 1 |
|  | Medium | M-TB | **<0.05** |
|  |  | M-R | **<0.05** |
|  |  | TB-R | 1 |
|  | High | M-TB | **<0.05** |
|  |  | M-R | **<0.05** |
|  |  | TB-R | 1 |

### S1.3 Figures

**Fig. S1:**
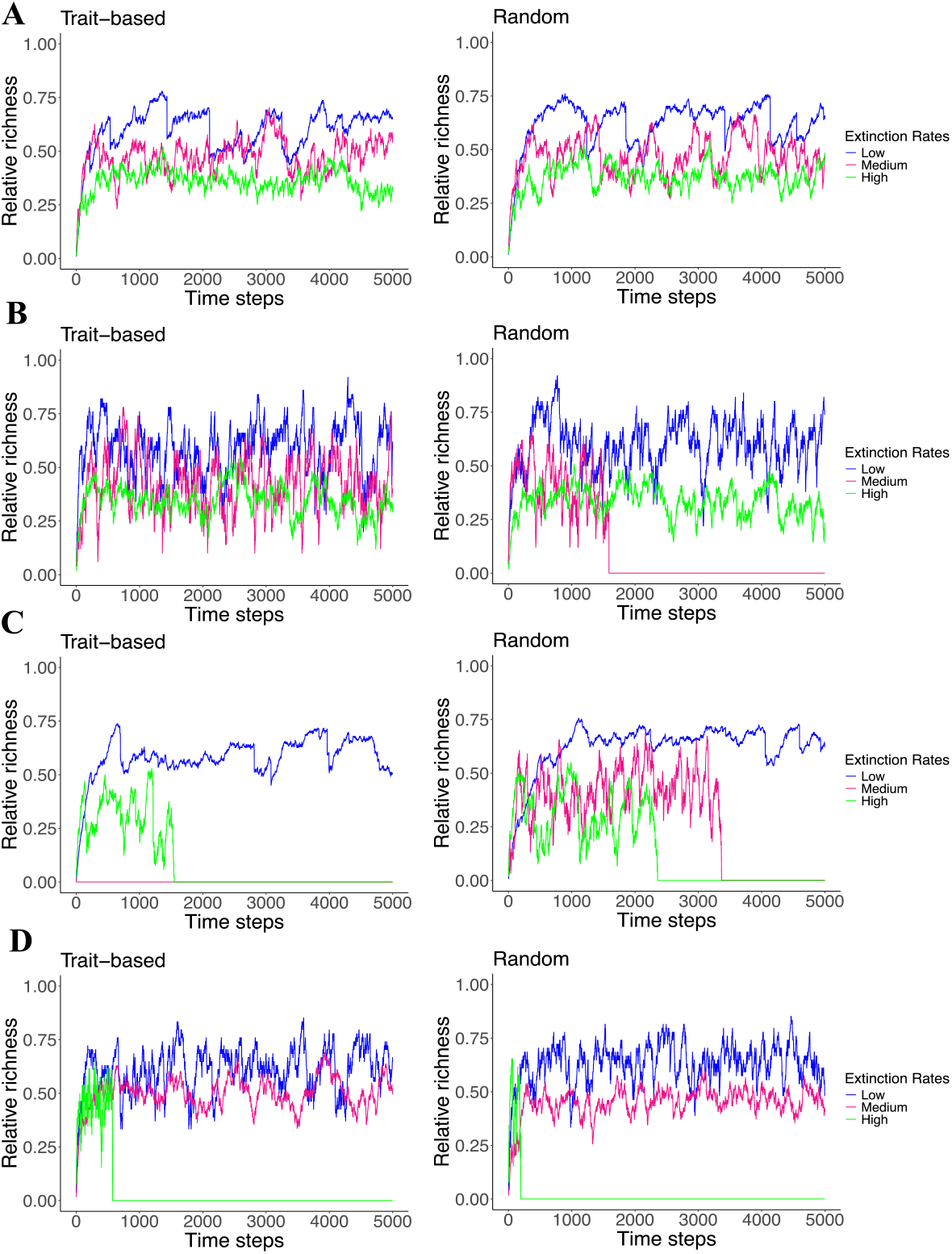
Richness of the island relative to total richness in the mainland for four different networks (**A-D**) in trait-based and random extinctions scenarios.

**Fig. S2:**
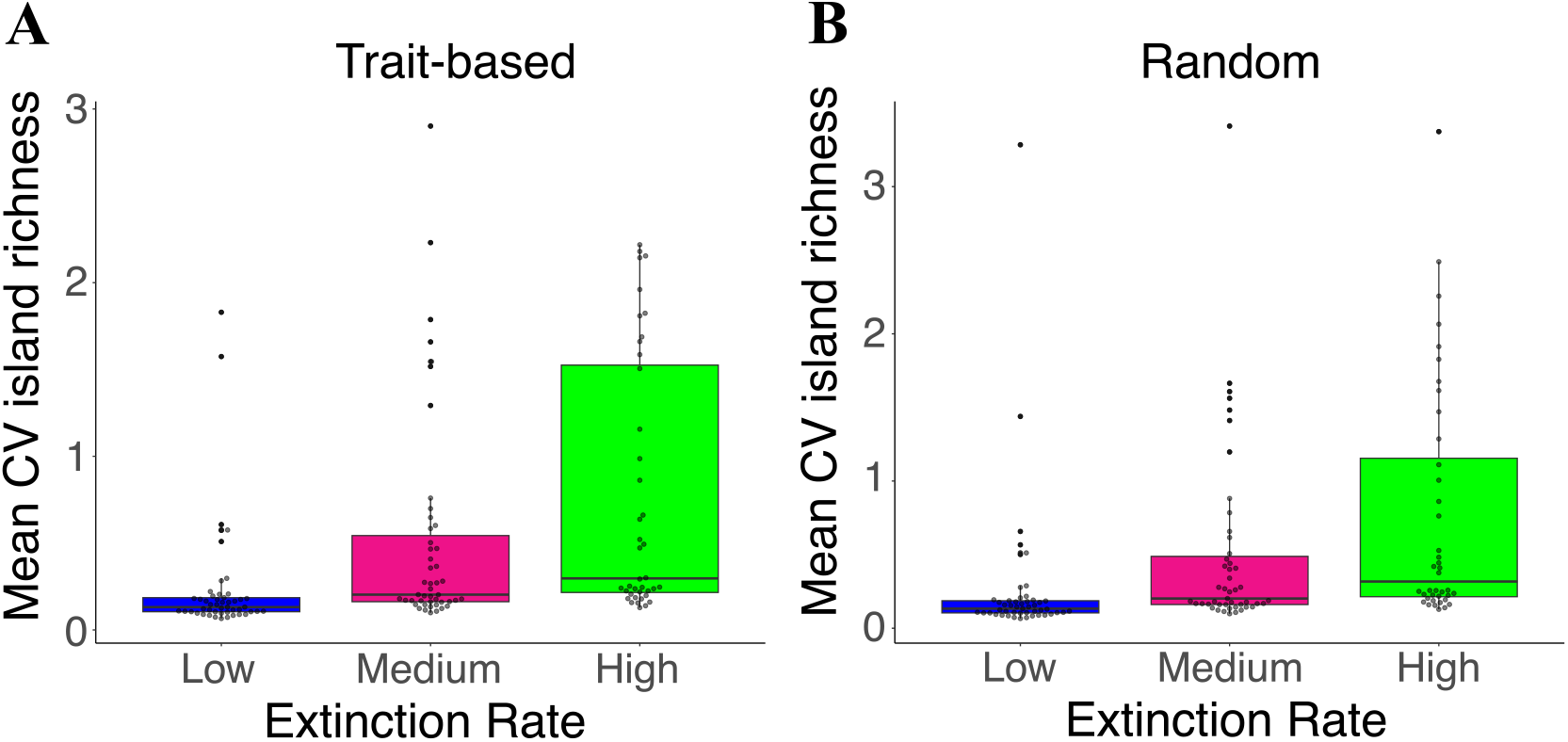
Mean CV of island richness communities across 1000 simulations at different extinction rates. Each point represents a network. **A)** Trait-based extinction scenario and **B)** random extinction scenario.

**Fig. S3:**
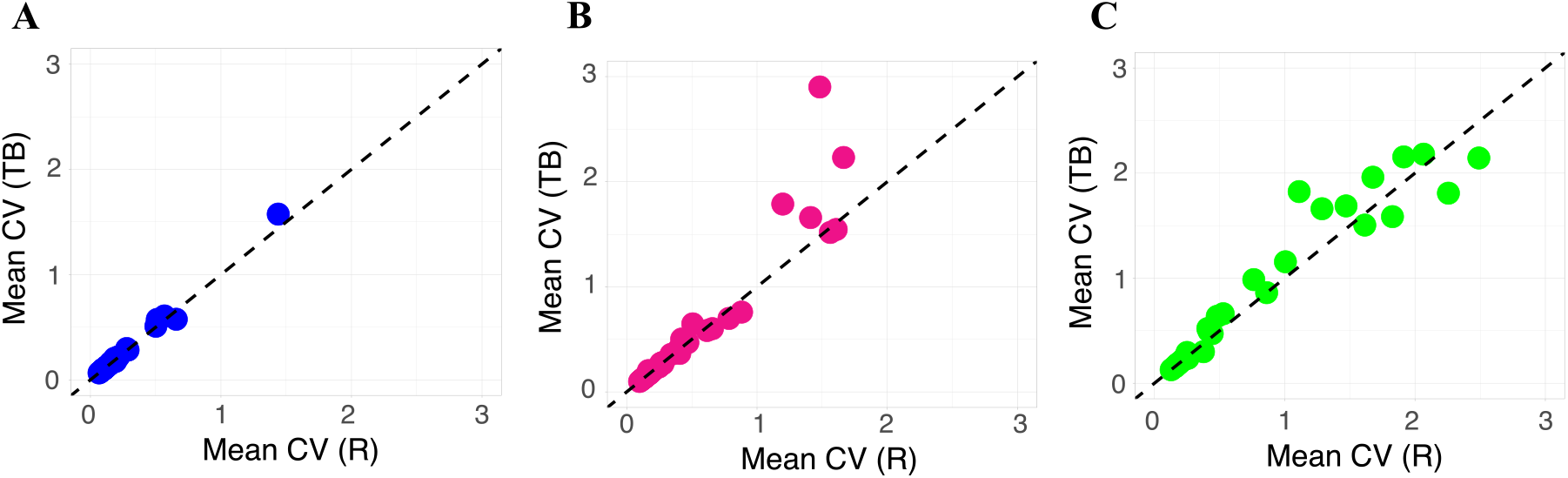
Correlation between the mean CV values of island richness in trait-based (TB) and random (R) scenarios under **A)** low, **B)** medium, and **C)** high extinction rates. Each data point represents the mean CV of a network calculated from 1000 simulations.

**Fig. S4:**
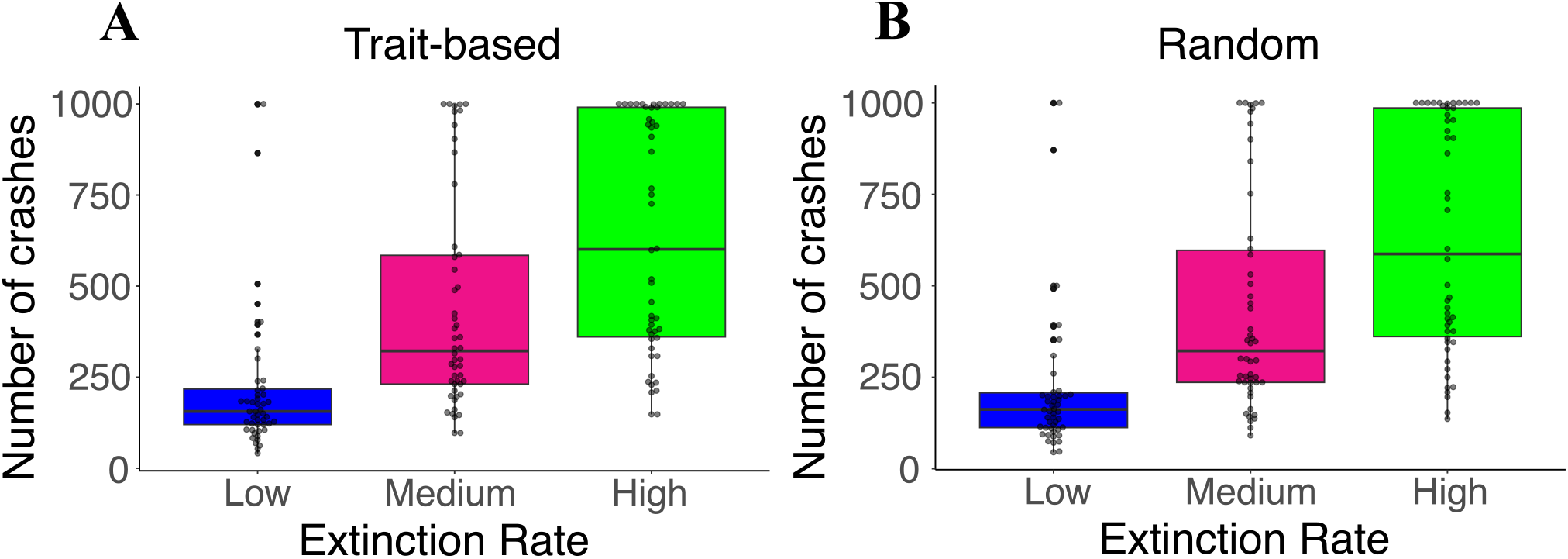
Number of crashes in island communities across 1000 simulations at different extinction rates. Each point represents a network. **A)** Trait-based extinction scenario and **B)** Random extinction scenario.

**Fig. S5:**
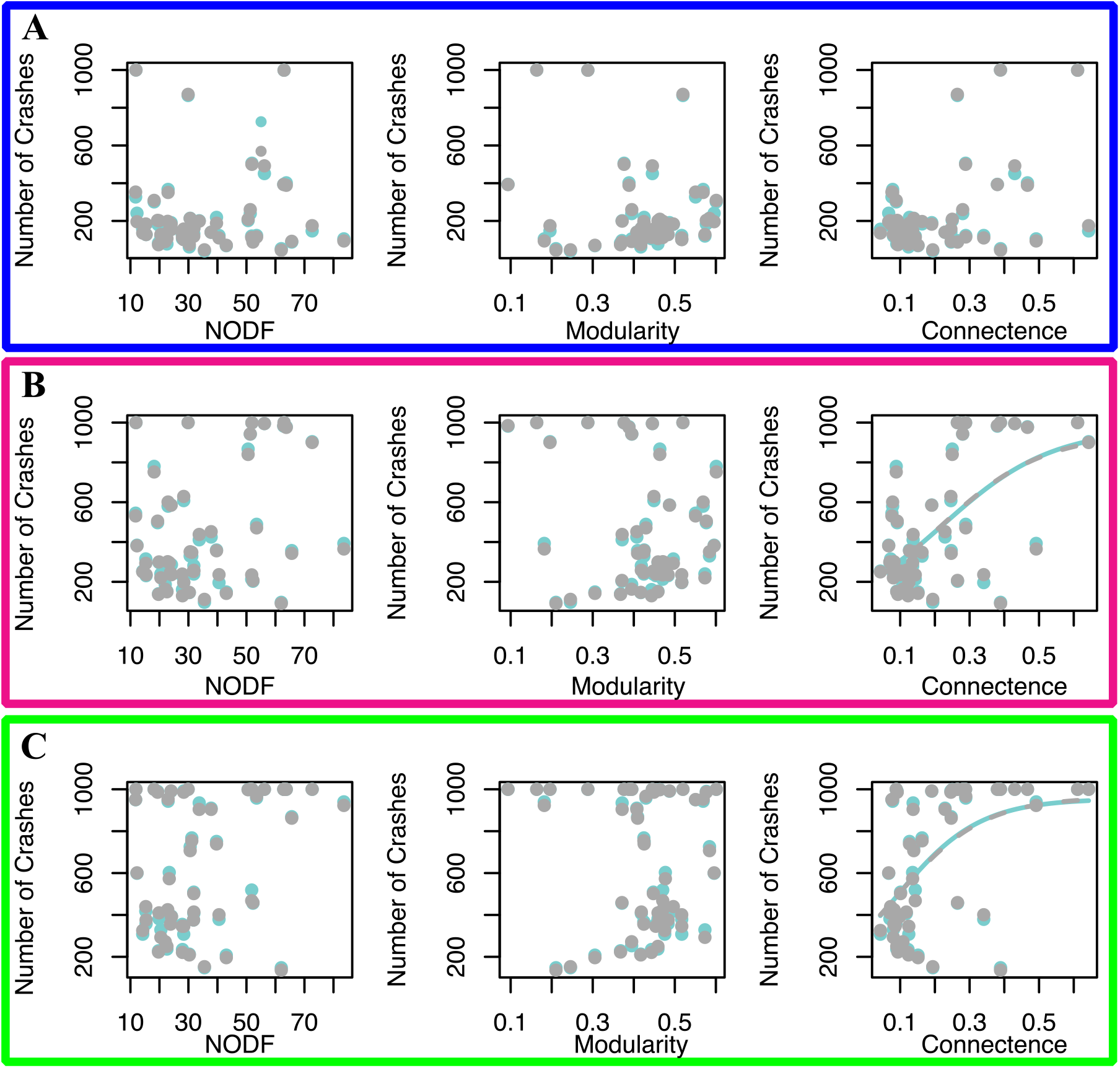
We fitted a logistic regression model to assess how the structure of mainland networks impacts the number of island community collapses across three extinction rates: **A)** Low (blue), **B)** Medium (pink), and **C)** High (green).

**Fig. S6:**
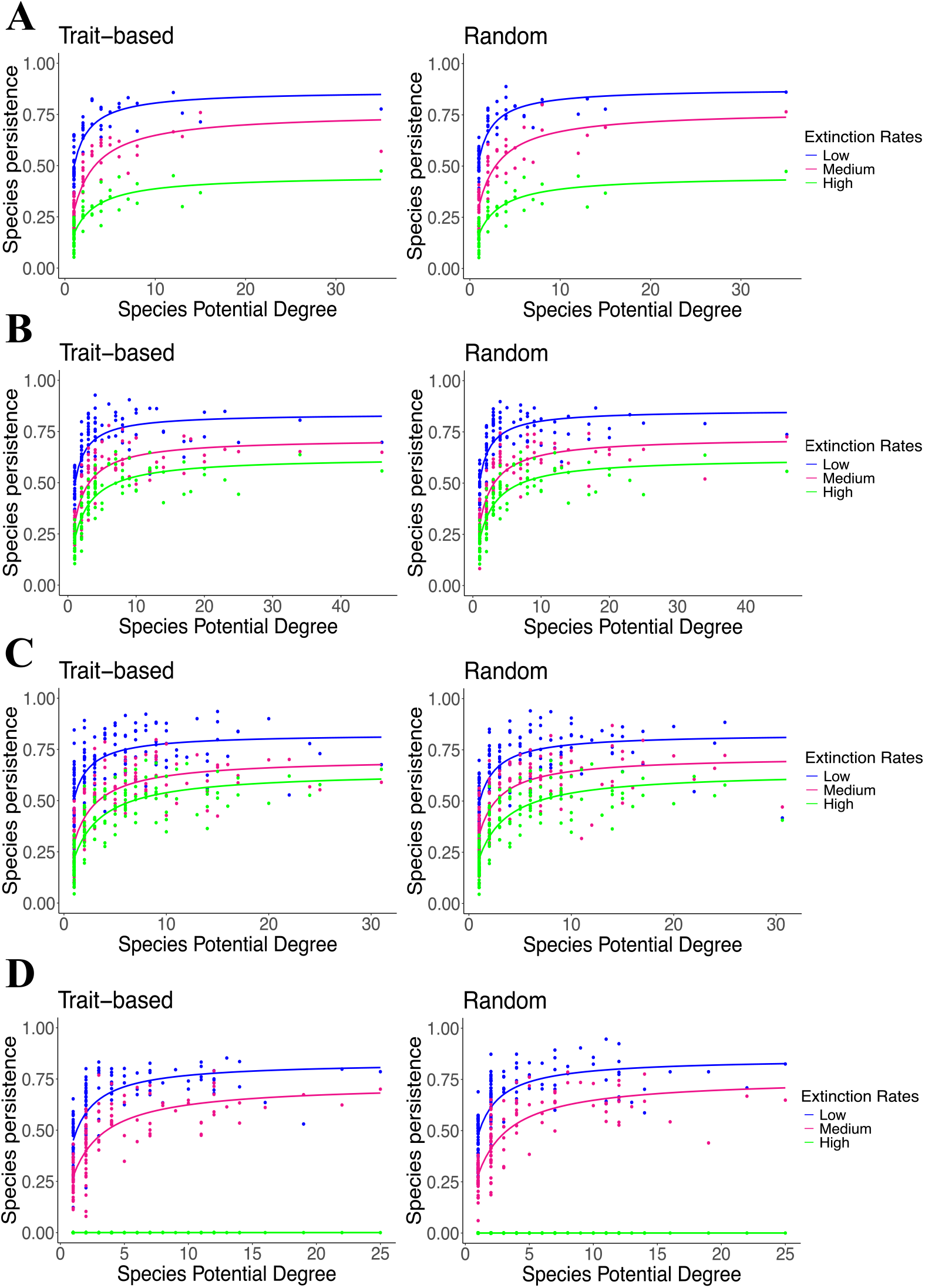
Species persistence on the island is related to the number of interactions species have on the mainland for four different networks (**A-D**) in trait-based and random extinction scenarios. Maximum persistence (= 1) means that the species was present on the island throughout all 5000 time steps in our simulation.

